# Large haplotypes highlight a complex age structure within the maize pan-genome

**DOI:** 10.1101/2022.02.22.481510

**Authors:** Jianing Liu, R. Kelly Dawe

## Abstract

The genomes of maize and other eukaryotes contain stable haplotypes in regions of low recombination. These regions, including centromeres, long heterochromatic blocks and rDNA arrays have been difficult to analyze with respect to their diversity and origin. Greatly improved genome assemblies are now available that enable comparative genomics over these and other non-genic spaces. Using 26 complete maize genomes, we developed methods to align intergenic sequences while excluding genes and regulatory regions. The centromere haplotypes (cenhaps) extend for megabases on either side of the functional centromere regions and appear as evolutionary strata, with haplotype divergence/coalescence times dating as far back as 450 thousand years ago (kya). Application of the same methods to other low recombination regions (heterochromatic knobs and rDNA) and all intergenic spaces revealed that deep coalescence times are ubiquitous across the maize pan-genome. Divergence estimates vary over a broad time scale with peaks at ∼300 kya and 16 kya, reflecting a complex history of gene flow among diverging populations and changes in population size associated with domestication. Cenhaps and other long haplotypes provide vivid displays of this ancient diversity.

## INTRODUCTION

The origins of maize can be traced to stands of teosinte, a tall grass with many small ears that is native to Mexico (Doebley 2004). Teosintes are divided into four species, one of which is *Zea mays*, which is divided into three subspecies: *Zea mays mays* (cultivated maize), *Zea mays parviglumis*, and *Zea mays mexicana*. Genetic data suggest that maize originated from populations of *parviglumis* (Matsuoka et al. 2002) with lesser contributions from *mexicana* (Ross-Ibarra et al. 2009; Hufford et al. 2013; Calfee et al. 2021). Through selection of desirable traits such as fewer stems and larger ears with exposed kernels, early inhabitants transformed teosinte into a domesticated crop as early as 8,700 years ago (Piperno et al. 2009). Domesticated maize was then transported northward through the desert and into Canada, south through the Andes, and east to the islands of the Caribbean where at each point it was cultivated as local landraces (Ross-Ibarra et al. 2009; van Heerwaarden et al. 2011; Hufford et al. 2013). Although the process of domestication caused a loss of genetic diversity relative to ancestral teosinte (Tenaillon et al. 2004; Wang et al. 2017), the remaining variation has been sufficient to sustain decades of continuous improvement by breeders (Andorf et al. 2019; Haberer et al. 2020; Hufford et al. 2021).

Molecular data show that there remains extraordinary variation in genome size, gene content, methylation status and repeat composition among maize lines (Chia et al. 2012; Sun et al. 2018; Haberer et al. 2020; Hufford et al. 2021). A large share of the diversity is a result of transposon insertion over the past three million years, which inflated genome size by two to five fold (Sanmiguel and Bennetzen 1998) and altered gene spacing and arrangement (Fu and Dooner 2002; Brunner et al. 2005). Transposable elements make up ∼83% of any single assembled maize genome (Hufford et al. 2021) and exhibit extreme polymorphism among maize lines. Dramatic examples are the two divergent haplotypes of the *bronze1* region (Fu and Dooner 2002), where there is an almost total absence of homology in the intergenic spaces: among 23 transposons annotated over ∼180 kb of combined sequence, only one transposon is conserved. Maize also contains several classes of tandem repeat arrays including those with *CentC*, a centromere repeat, and two interspersed repeats known as knob180 and TR-1 defining heterochromatic domains called knobs (Liu et al. 2020). The lengths of tandem repeat arrays vary over orders of magnitude and are remarkably polymorphic among lines (Albert et al. 2010). Much of this structural diversity is presumed to be a product of ancient haplotype divergence, predating speciation. Persistence of ancient haplotype diversity is attributable to a slow rate of genetic drift in large populations (Hilton and Gaut 1998; Clark et al. 2004) or gene flow across species and subspecies (Ross-Ibarra et al. 2009).

Highly repetitive, structurally divergent regions display reduced levels of recombination, allowing identification of large, often megabase-scale haplotypes that have persisted and diverged over thousands to a few hundred thousand years. Well known examples are the evolutionary strata on the human X chromosome that reflect the timing of major structural rearrangements (Lahn and Page 1999). Outside of sex chromosomes and other special cases, recombination is lowest around centromeres where it trends toward zero (Shi et al. 2010; Nambiar and Smith 2016). In human, complete genome assemblies have shown that centromeres occur in large haplotypes (called cenhaps) that harbor rich and previously unknown genetic diversity (Langley et al. 2019; Altemose et al. 2021). Here we sought to thoroughly describe the structure and evolution of maize haplotype blocks around centromeres, knobs and the nucleolus organizer region (NOR). We took advantage of 26 complete genomes from diverse maize inbreds (Hufford et al. 2021) and developed custom tools to carry out accurate alignment over TE-rich regions and repeat arrays. We found that maize centromeres, knobs and NOR regions exhibit striking haplotype diversity including segregating variants that exhibit a broad range of coalescence times predating the early domestication of maize. We interpret the distribution of haplotype coalescence times in the context of historical effective population size and gene flow among *Zea* subspecies.

## RESULTS

### Whole genome alignment over intergenic spaces

To investigate haplotype structure at genome scale, we analyzed 26 high quality genomes from the maize Nested Association Mapping (NAM) population (Hufford et al. 2021), a rich collection including temperate lines, tropical lines, sweet corn and popcorn (McMullen et al. 2009). We implemented the longest increasing subsequence (LIS) algorithm (Rani and Rajpoot 2016; Abouelhoda and Ohlebusch 2005) in a two-step chaining procedure (Sup Fig 1). This method resolved mis-alignment errors and effectively captured rearranged segments (Sup Fig 2). The LIS method also revealed ∼19 million structural variations (Sup Fig 3); a greatly expanded list (relative to an earlier database of ∼0.79 million; (Hufford et al. 2021)) which is available for use in mapping and association studies (Sup Fig 4, Sup Table 1 and 2). By comparing genomes in an all-by-all manner and retaining the unique portions contributed by each, we estimated the total (genic and intergenic) pan-genome size to be ∼7.9 Gb (Sup Fig 5A) which is ∼3.7 times larger than the average assembled size of any single genome (Hufford et al. 2021). Only ∼4.9% of the total pangenome (0.38 Gb) is conserved among all lines with the remaining 7.5 Gb segregating among lines at various frequencies (Sup Fig 5B).

### Ancient haplotypes in centromeric regions

Alignments revealed megabase-scale cenhaps on seven chromosomes, including a ∼5 Mb region on chromosome 2, a ∼10 Mb region on chromosome 3, a ∼7 Mb region on chromosome 5, a ∼10 Mb region on chromosome 7, a ∼10 Mb region on chromosome 8, a ∼7 Mb region on chromosome 9 and a ∼8 Mb region on chromosome 10 (Sup Fig 6, 7). The segregating haplotype variants include the functional centromere regions defined by the presence of Centromeric Histone H3 (CENH3) (Wang et al. 2021; Hufford et al. 2021) and extend well into flanking pericentromeric sequences. For example, five inbreds CML52, HP301, IL14H, Mo18W, and P39 share a cenhap on chromosome 8 that is clearly distinct from the cenhap variant seen in all other inbreds (Fig 1A, B). There is extreme variation in TE distribution between these two chromosome 8 cenhaps (Fig 1C), mirroring the TE polymorphism previously described at the *bronze1* locus (Fu and Dooner 2002). Most maize centromeric regions also contain long arrays of a ∼156 bp tandem repeat known as CentC (Gent et al. 2017). On chromosome 8, the CentC array within the major cenhap group (the B73 type) and alternate type (the P39 group) show no collinear homology (Fig. 1D), suggesting a complete turnover of repeat arrays. All-by-all clustering of the chromosome 8 data further supports the existence of two deeply divergent haplotype groups (Fig 1E). Similarly divergent CentC arrays are observed on chromosomes 2 and 3 (Sup Fig 8, 9).

**Figure 1.**
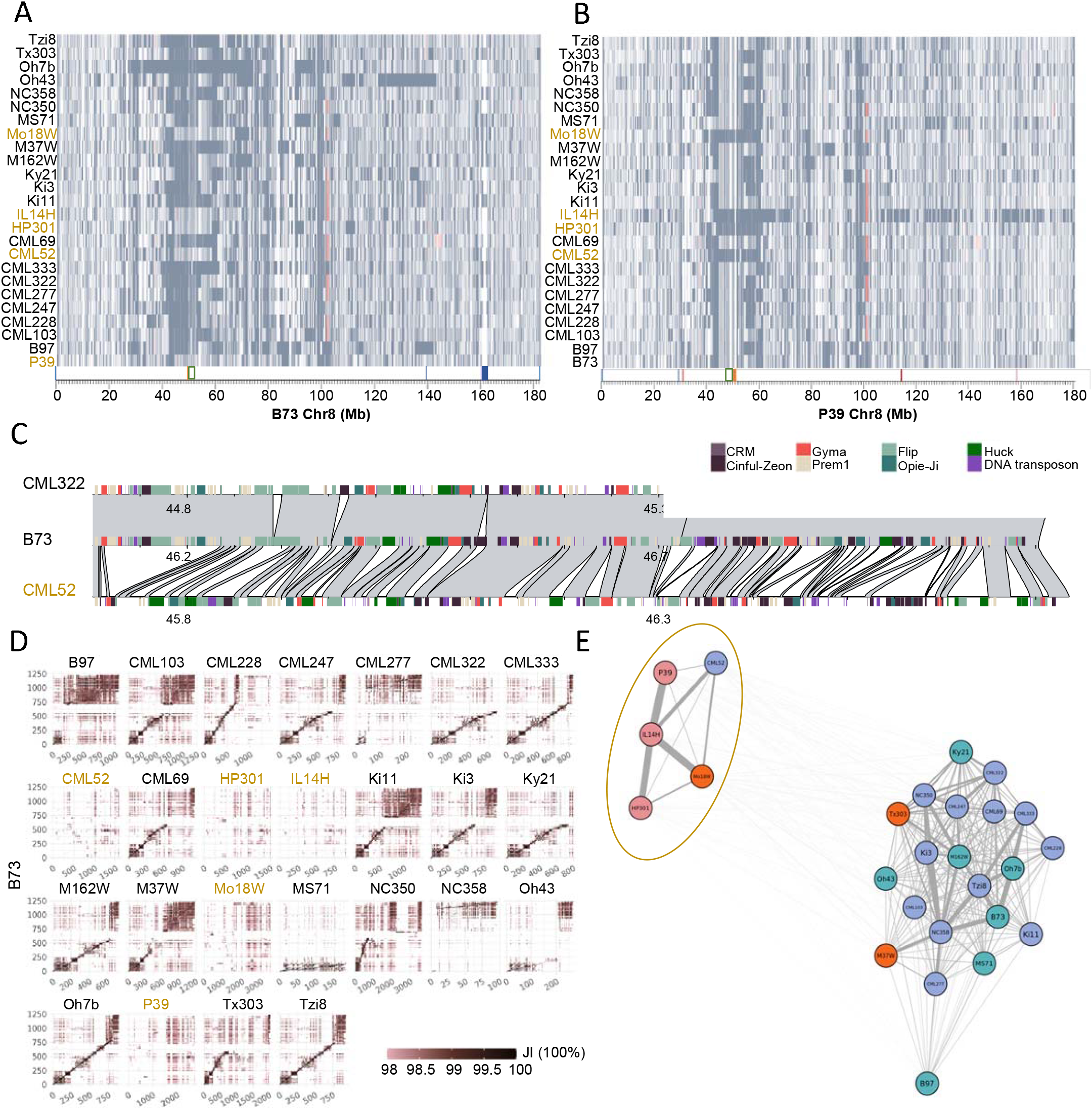
Cenhaps on chromosome 8. **A)** Alignments between NAM lines and B73 over chromosome 8. Syntenic aligned regions (grey) and inverted segments (red) are shown. CENH3 Chip-seq regions (green box) CentC (orange), knob180 (blue), and TR-1 knob (red) are highlighted on the x axis. **B)** Alignments between NAM lines and P39 over chromosome 8. Annotation is the same as A. **C)** Pairwise alignments and TE comparisons between CML322, CML52 and B73 over a ∼1 Mb region of chromosome 8. Major TE families are noted in colors. This region does not include the functional centromere domain. **D)** Pairwise alignments between NAM lines and B73 over CentC arrays on chromosome 8. X and y axes show CentC monomers, and color intensity reflects the Jaccard index between each monomer pair. The non-B73 haplotypes are shaded in olive. **E**) Clustering of CentC arrays on centromere 8. The colors over inbred names indicate varieties of corn: northern flint (pink), temperate (blue), mixed (red), and tropical (green).

To compare the age of maize cenhaps, we identified SNPs in the aligned regions for each chromosome of the B73 inbred and calculated the times of divergence assuming a mutation rate of 3.3 × 10^−8^ substitutions per site per year (Clark et al. 2005) (Fig. 2, Sup Fig 10). Before doing so, we masked all annotated genes and unmethylated (potential regulatory) regions under the assumption that intergenic sequences would be less likely to be constrained by selection (Lynch et al. 2016; Monroe et al. 2022). The divergence dating confirmed that cenhaps on seven chromosomes fall into two haplogroups with divergence times ranging from ∼450-130 mya. Cenhaps often contain segments of lesser age, suggesting occasional recombination. For example, the cenhaps on chromosome 10 include sections with estimated divergence times (from B73) of 450 kya intermixed with 130 kya segments (Fig 2A).

**Figure 2.**
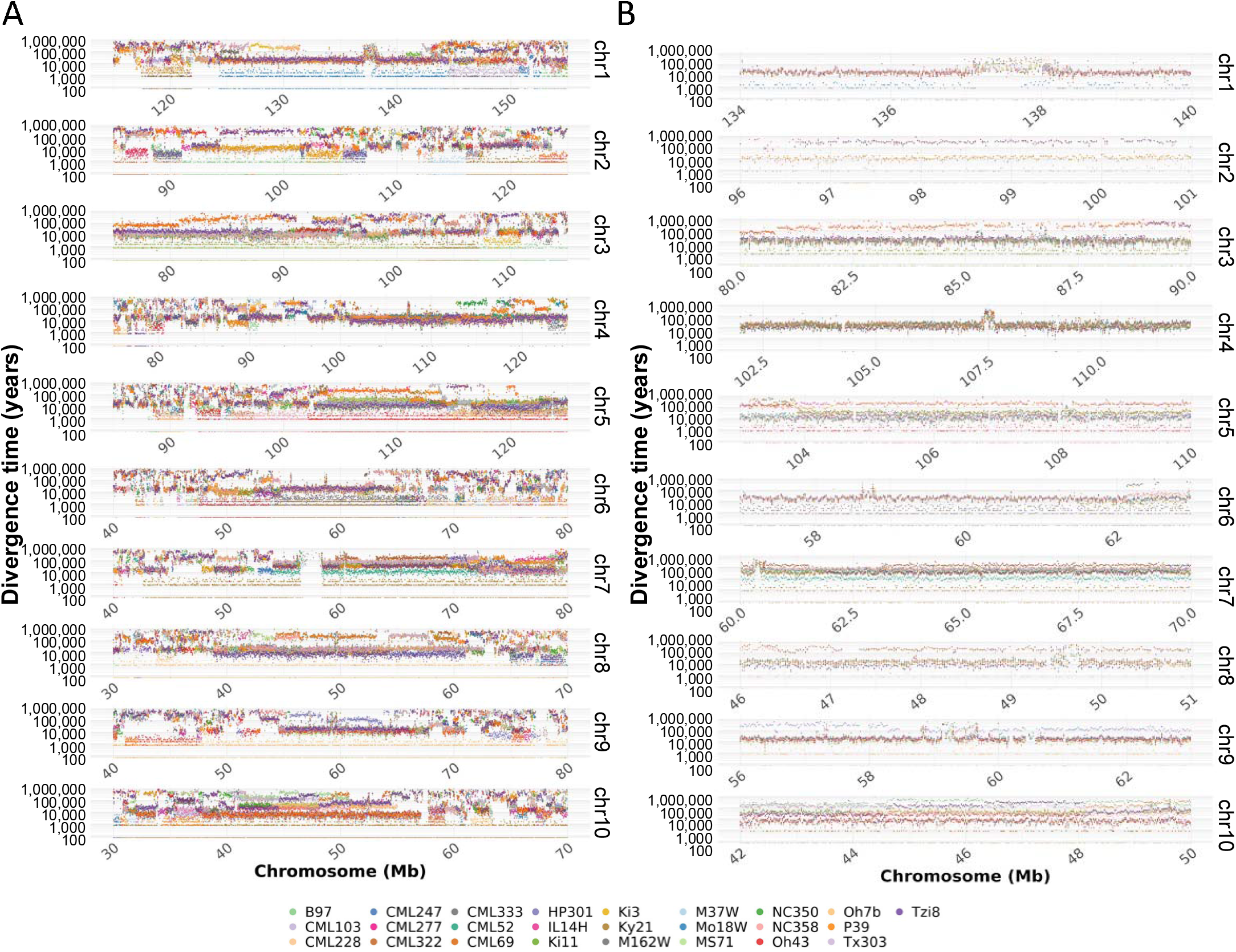
Divergence times of cenhaps. **A)** Divergence times between NAM lines and B73 over selected pericentromeric regions. Each dot represents an estimated divergence time over a 20 kb window. The cenhap regions are highlighted in boxes. **B**) Zoomed-in view of cenhaps outlined in A. The divergence times over CentC regions (orange bars) are not reliable because of non-syntenic alignment of centromeric retrotransposons (CRMs) within the arrays.

The times of cenhap divergence coincide with the estimated origin of the *Zea* lineage (∼100-300 kya; (Ross-Ibarra et al. 2009)). We tested whether the major cenhaps are present in teosintes by aligning Illumina data from 67 teosinte accessions including *Zea mays* ssp. *parviglumis*, ssp. *mexicana*, ssp. *huehuetenangensis* and the related species *Zea diploperennis* to B73. The presence and absence of major cenhap groups could be scored based on the density of aligned reads, although there were ambiguous cases where the centromeres were heterozygous and sequence coverage was low (Sup Fig 11). The data demonstrate that all major cenhap groups found in maize (chromosomes 2, 3, 5, 7, 8, 9, 10) and two additional cenhaps on chromosome 1 and 6 also occur as segregating polymorphism in *parviglumis*. Among the four *mexicana* accessions analyzed, there were alternate cenhaps on at least five chromosomes (chromosome 3, 4, 5, 7, 10).

### Ancient haplotypes in repeat arrays of knobs and NOR

*Zea* species contain many heterochromatic knobs with megabase-scale arrays of tandem repeats known as knob180 and TR-1. Knobs are found in mid-arm positions and are maintained by a meiotic drive mechanism (Dawe et al. 2018; Swentowsky et al. 2020). Recombination is suppressed within and around knobs (Ghaffari et al. 2013). Analysis of nine large knobs in the NAM lines (Albert et al. 2010; Hufford et al. 2021) demonstrated that two (6L, 8L) fall into two clusters that have diverged for over 200 kya (Sup Fig 12 and 13). The knob on the short arm of chromosome 9 (9S) separates into three clusters, where two diverged from B73 over 300 kya (Fig. 3A,B). In contrast, six knobs have divergence times of less than 100 kya consistent with a more recent emergence, presumably as an outcome of meiotic drive. We also analyzed the nucleolus organizer region (NOR) which contains megabase-scale arrays of rDNA (Fig 3C). All-by-all alignment of the 6S knob linked to the NOR indicated three distinct clusters with progressive divergence times of 100, 220 and 300 kya (Fig 3D). The analysis also corroborates the recent report of a ∼3 Mb insertion of non-rDNA sequence with homology to *Tripsacum dactyloides* (the sister genus to *Zea*; Sup Fig 14) within the most common maize NOR haplotype (Huang et al. 2021), but minor NOR haplotypes do not include this insertion.

**Figure 3.**
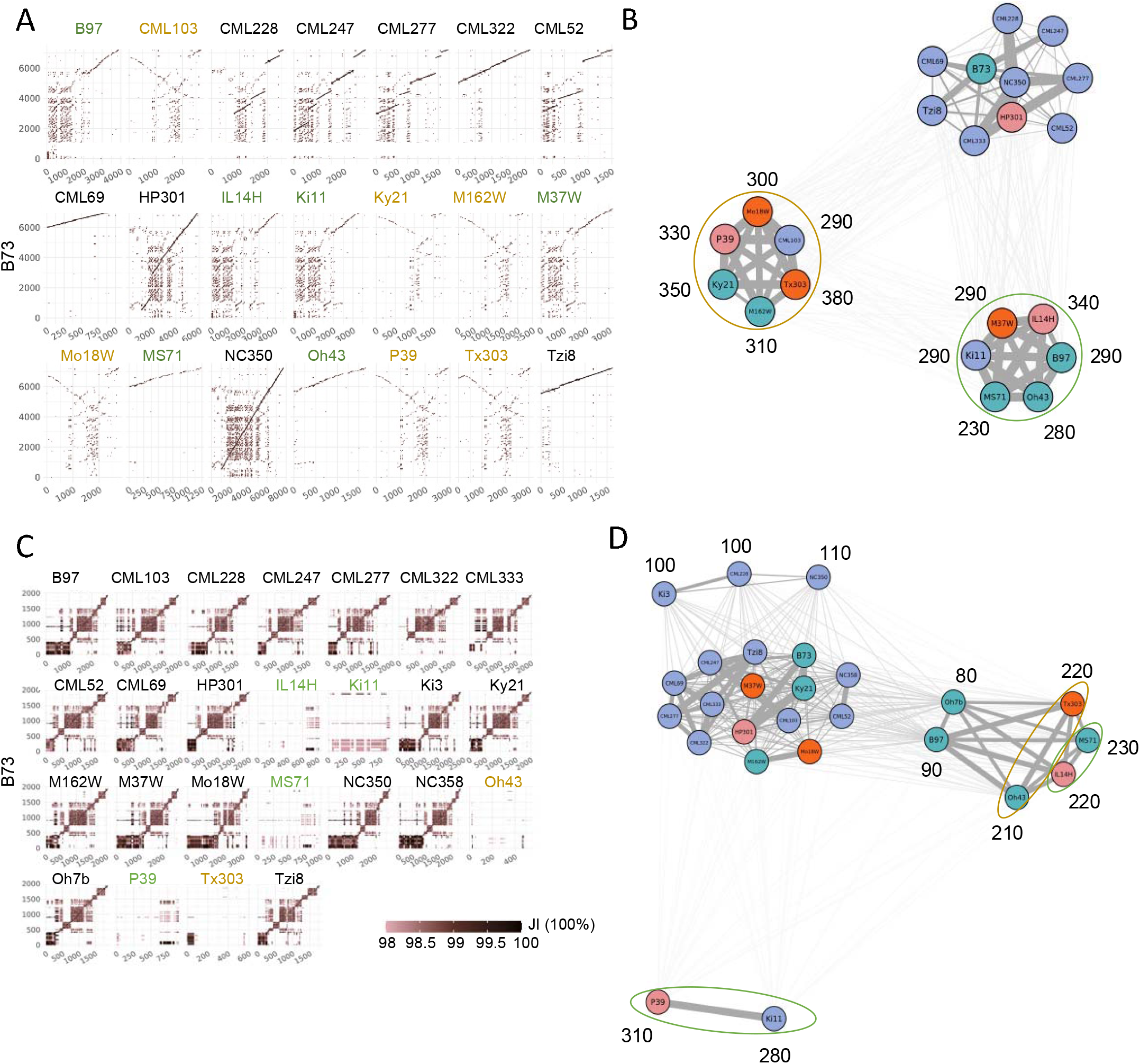
Ancient haplotypes in the 9S knob and NOR. **A)** Dot-matrix alignments between 21 NAM lines and B73 over the knob180 array on chromosome 9S (only 21 have the 9S knob). X and y axes show knob180 monomers, and color intensity reflects the Jaccard index between each monomer pair. **B**) Clustering of 9S knobs based on repeat array alignments. Divergence times (kya) were calculated using syntenic SNPs in TEs that lie within the arrays. **C**) Dot-matrix alignments between 25 NAM lines and B73 over the NOR. X and y axes show 18S rDNA monomers, and color intensity reflects the Jaccard index between each monomer pair. **D**) Clustering of the 6S knob180 array that is about 10 Mb from NOR on chromosome 6. The TEs in the knob can be dated using syntenic SNPs but there has been some recombination between NOR and 6S knob. In A and C, the non-B73 haplotypes are indicated in olive or green, and the corresponding clusters in B and D are circled in matching colors. In B and D, the colors over individual inbred names indicate varieties of corn: northern flint (pink), temperate (blue), mixed (red), and tropical (green).

### A pangenome-wide burst of diversity at 30-10 kya

Our documentation of maize cenhaps (Fig 2) reveal not only extreme cases of 300-100 kya of divergence between non-recombining variants, but also a significant amount of divergence in the 30-10 kya range. These divergence times were estimated against a single reference genome. To gain a broader perspective we also performed hierarchical clustering using SNPs within cenhaps to interpret relative divergence in an all-by-all manner. The results confirm the deep separation of major haplotype groups and demonstrate that the majority of modern cenhap diversity evolved in the recent past (Fig 4A and Sup Fig 15).

**Figure 4.**
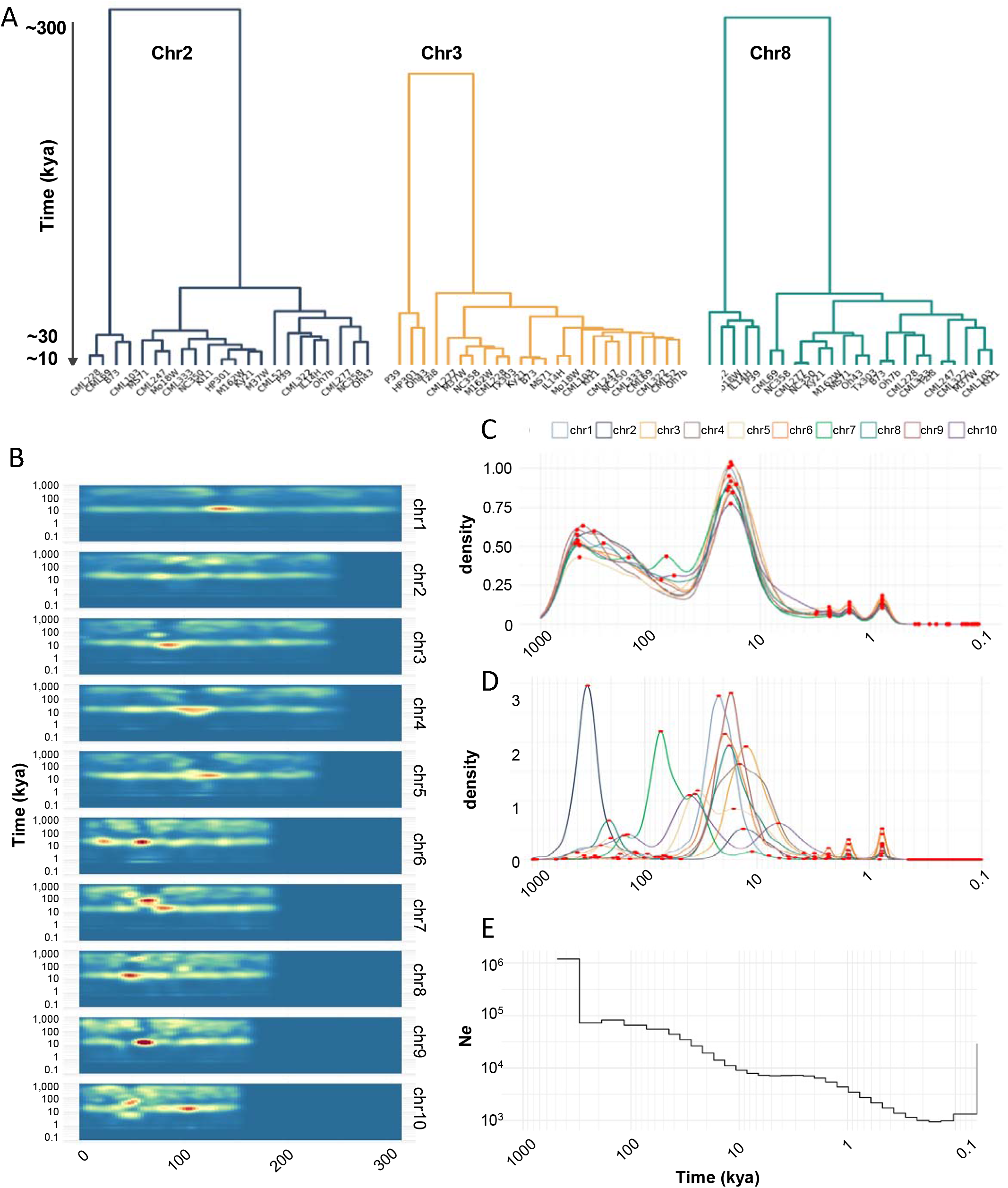
Whole genome age distributions. **A)** Hierarchical clustering of cenhaps from three chromosomes showing that most cenhap diversity dates to 30-10 kya. **B)** Divergence times of intergenic spaces estimated from 20 kb windows represented in a density plot, where the highest density is indicated in red. **C)** Density plot of all divergence times displayed in B. The x axis shows probability densities, where the total area under the curve is 1. Local maxima are highlighted as red dots. The peaks on the tail of the distribution (at ∼2.4, 1.6, and 0.8 kya) represent windows with 3, 2, or 1 SNP. **D**) Density plot of cenhap divergence times only, annotated as in C. **E)** Effective population size (N*e*) of maize over the past 0.5

To determine whether these trends are unique to cenhaps, we plotted molecular age densities across all chromosomes of all NAM lines using B73 as a reference. This effort was designed to focus on the tens of thousands of smaller haplotypes between genes. The intergenic spaces in maize are about ∼50 kb on average (∼40,000 genes in ∼2.2 Gb genomes) and most recombination occurs within genes and areas with unmethylated DNA (Liu et al. 2009; Choi et al. 2018), which were omitted from our alignments. We chose to estimate divergence times using 20 kb sliding windows, but observed the same patterns with window sizes ranging from 10-100 kb (Sup Fig 16). The results clearly show that molecular age distributions are not uniform, with multiple bands of coalescence times in the 500-100 kya range and a rich layer of diversity dating to 30-10 kya (Fig 4B). These general trends are more apparent when the genome-wide divergence data are displayed as a single age density plot (Fig 4C, Table 1). In very broad terms, the age distributions can be seen as bimodal, with about 29.4% of the total diversity originating between 500-100 kya, and ∼32.7% of the diversity originating in a burst-like pattern between 30-10 kya (Fig 4C, Table 1). We observed similar trends when the analysis was performed on cenhaps alone (Fig. 4D), indicating that cenhaps, while large and conspicuous, are not otherwise unusual.

**Table 1.** Percent of pan-genome that diverged during different time intervals.

| Time interval | # Years included | Percent of pan-genome diverged during interval <sup>1</sup> | # SNPs in interval |
| --- | --- | --- | --- |
| 1,000 - 500 kya | 500,000 | 7.5 | 65,613,821 |
| 500 - 100 kya | 400,000 | 29.4 | 92,393,623 |
| 100 - 30 kya | 70,000 | 13.7 | 15,955,235 |
| 30 - 10 kya | 20,000 | 32.7 | 15,235,449 |
| 10 - 0.1 kya | 9,900 | 11.1 | 1,487,184 |

<sup>1</sup> The pan-genome is defined as the sequence present in the 25 NAM founder inbreds. The genomes from all 25 lines were aligned in 20 kb windows to the B73 reference; this column shows the percent of windows with the indicated divergence time from B73. Areas that are identical by descent (defined as less than 0.1 kya divergence) are not included.

The peak of coalescence times in the 30-10 kya timeframe is notable for its steep flanks on both sides. On the left side of the peak, the transition from relatively few alleles dating to 100-30 kya to an abundance at 30-10 kya can be explained as a natural outcome of genetic drift in populations with finite effective population sizes (*Ne*) (Wright 1949; Nordborg 2004). On average, two alleles will coalesce to their most recent common ancestor after 2*Ne* generations. Three prior estimates of historical *Ne* suggest a minimum *Ne* for maize of ∼6,000 (Beissinger et al. 2016), a decreasing *Ne* starting at ∼50,000 and declining to ∼1000 during domestication (Wang et al. 2017), and a decreasing *Ne* starting at about ∼100,000 and declining to ∼10,000 during domestication (Tittes et al. 2021). All prior analyses were carried out using short-read alignments. We carried out a fourth analysis of historic *Ne* using our whole genome alignments of 26 genomes (Fig 4E). The results suggest that *Ne* dropped from a high of ∼90,000 to ∼1000 during domestication (the sudden change in *Ne* at ∼130 kya is probably spurious (Patton et al. 2019)). Interestingly, we infer from our data that the population stabilized at about 8,000 individuals for several thousand years (in the period 6-2 kya). Applying 2*Ne* to this stable population size predicts an average coalescence time of ∼16 kya, which matches the observed peak in estimated allele age at ∼16-17 kya (Sup Fig 16). Relatively little haplotype diversity dates directly to the domestication period when effective population sizes were at their lowest. Only about 11.1% of the 20 kb windows (Table 1) and 25 of the 260 cenhaps diverged from each other in the recent <10 kya past.

## DISCUSSION

Here we describe an approach for inferring evolutionary history based on SNP diversity within intergenic spaces. Repeat-rich, highly heterochromatic plant genomes such as maize include megabase-scale stable haplotypes in areas of low recombination, including cenhaps, knobs and the NOR. We show that analyses of coalescence times for haplotype variants can elucidate complicated population dynamics over the evolutionary history of extant species. Genome-wide comparisons revealed bands of varying age densities over a broadly bimodal trendline (Fig. 4B,C). Approximately 29.4% of the diversity in the maize pan-genome is ancient, originating between 500-100 kya (Table 1). These regions are structurally very different, with long regions of differential TE insertions and relatively little alignable sequence (Fig. 1C). In cenhaps there are extreme differences in CentC arrays, to the point that in some cases there is no evident collinear homology (Fig. 1D). Another ∼32.7% of the diversity emerged in a burst-like pattern 30-10 kya. This period coincides with major changes in earth’s climate (Leyden et al. 2013; Williams 2003) and the arrival of humans to the Americas (Ardelean et al. 2020; Becerra-Valdivia and Higham 2020). It is possible that some of the recent diversity accumulated as a result of demographic changes associated with these environmental events. However, the inferred reduction in population size to about 8000 during the 6-2 kya period is sufficient to explain the peak in coalescence times dating to 30-10 kya.

The evolutionary trendline of the maize pan-genome, with a significant amount of ancient diversity intermingled with a concentration of coalescence times at 30-10 kya, is clearly evident in the diversity of large haplotypes around the genome. Authors of a previous study argued that the apparent overabundance of recently evolved centromere haplotypes might be a direct outcome of domestication and selection (Schneider et al. 2016). Our analyses indicate that cenhap coalescence time distributions closely resemble and indeed accentuate genome-wide patterns of haplotype diversity (Fig. 4C,D). It has also been suggested that centromeres might evolve by meiotic drive, a process that could cause episodic sweeps of some cenhaps over others and reduce overall cenhap diversity (Malik 2009). While centromere drive does occur in some species and lineages, it is likely rare (Lampson and Black 2017; Finseth et al. 2021) and drive-based sweeps are not evident in the maize cenhap data. The persistence of at least two ancient cenhaps for seven of the ten maize centromeres (and two of the remaining centromeres in teosinte) suggests that centromere drive has not had a major impact in maize over the last ∼100 kya.

Given the small effective population sizes during domestication (Fig 4E), we might expect that relatively little ancient polymorphism would have survived into modern maize, yet extensive ancient diversity persists (Fig. 4C, Table 1). A likely explanation for the high levels of ancient polymorphism in maize is admixture among species and subspecies populations (Ross-Ibarra et al. 2009; Hufford et al. 2013). Maize is thought to have been domesticated from *Zea mays* ssp. *parviglumis* with concurrent or subsequent intercrossing with ssp. *mexicana* (Hufford et al. 2013; van Heerwaarden et al. 2011). Subspecies *parviglumis* has maintained consistently large effective population sizes over the last 10 kya (Wang et al. 2017) and contains substantially more genetic diversity than modern maize (Hufford et al. 2012; Beissinger et al. 2016). The Mesoamerican genus *Zea*, including *Z. mays, Z. luxurians, Z. diploperennis, Z. nicaraguensis, and Z. perennis*, is estimated to have originated from ∼300 to ∼100 kya (Ross-Ibarra et al. 2009). Our divergence time analyses reveal a coalescence time peak dating to ∼300 kya that may represent the origin of *Zea* (Fig 4C). Additional complete genome assemblies from multiple *Zea* relatives will help to clarify these relationships and the extent of gene flow among species and subspecies.

## METHODS

### Genome alignment and structural variant characterization

Previous studies of the NAM lines have interpreted structural variation by aligning resequencing data to the primary B73 reference genome, using both short and long-reads (Hufford et al. 2021; Chia et al. 2012; Gore et al. 2009). However, this approach fails for insertions that are larger than the read length and preferentially recovers deletions. A partial solution to this problem is the merged alignment blocks method that was used to identify structural variants with whole-genome alignments across six European flint lines (Haberer et al. 2020). While this approach identifies simple insertions and deletions accurately, it performs poorly in regions where the reference and query differ by multiple insertions/deletions. Another approach to improve mapping and variant detection is AnchorWave, which relies on gene annotation to anchor reference and query (Song et al. 2022). Our method was designed for regions with few genes and high TE content. In these areas, non-syntenic alignments are frequently observed due to homology among similar transposons (Sup Fig 2). To remove the alignment noise, we identified syntenic regions by identifying Longest Increasing Subsequences (LIS) (Rani and Rajpoot 2016; Abouelhoda and Ohlebusch 2005) in a two-step procedure (Sup Fig 1 and 2).

The workflow consisted of three phases: 1) Perform pairwise whole-genome alignment, 2) Chain aligned segments with LIS in a two-step process, where the first step identified syntenic (collinear) segments, and the second step resolved locally rearranged regions, and 3) Characterize structural variants through identifying alignment gaps.

#### Alignment

Genome assembly and annotation files were obtained from MaizeGDB (https://maizegdb.org/NAM_project). Pairwise genome alignments were carried out with minimap2 (Li 2018) (v2.17) using parameters: -c -cx asm5 --no-kalloc --print-qname --cs=long. The alignments were then sorted according to the position in the reference sequence.

#### Chaining

A two-round chaining procedure was implemented to identify the longest set of anchors, where the first round identifies the optimal chain and the second round finds lower-scoring anchors to fill the gaps in the first chain (Sup Fig 1). During each round, we calculated the chaining score for individual anchors, and identified non-overlapping anchors in the global optimal path using the backtracking approach. The computation of chaining score differed between the two rounds, as the second was carried out to incorporate anchors of lower mapping quality.

The chaining score of anchor *i* in the first round was calculated as:*f*(*i*) = *max*{*f*(*j*) + *len*(*i*) * *log*10(*q*(*i*) + 0.001) − *gap*(*i, j*)/100}, *i* > *j* > 1, where len(i) and q(i) are respectively the length and the mapping quality of anchor i. Gap (i,j) is the distance between anchors i and j, which was computed as *abs* (*ix* − *jy*), and x and y are the start and end coordinates. After score calculation, the backtracking method was used by repeatedly finding the best predecessor of anchor i. After the first round, anchors identified in the optimal chain were combined with the remaining anchors larger than 15 kb and subjected to round two, where the score calculation of anchor i was modified from step1 as: *f*(*i*) = *max*{*f*(*j*) + *len*(*i*) − *gap*(*i, j*)/100}, *i* > *j* > 1.

#### Structural Variant characterization

In cases where two genomes differ by independent TE insertions in the same region, there can be unalignment gaps of different sizes. This special variant structure cannot be characterized by software developed to score small variants or variants with simple junctions. To accurately characterize structural changes in both reference and query genomes, we defined these variants as pairwise unaligned regions (Sup Fig 3B). Translocations and tandem duplications were inferred from alignment chains and orientation. True variants were further filtered with a 20 kb cutoff for inversions, 10 kb for tandem duplications, and 50 kb for translocations.

### Variant identification among NAM lines

To identify structural changes among 26 NAM lines, we performed 325 pairwise alignments for each chromosome and carried out chaining and SV characterization with the workflow described above. Chaining was conducted with script “chaining.py” and SV calling was accomplished with “sv_detect.py” (see WholeGenome-SV section in github). The number and size of variants, including un-alignments, tandem duplications, and inversions, were quantified for individual genomes and plotted with custom script using karyploteR. For each pair of unaligned regions, the region in the reference was defined as a deletion, and its counterpart in the query an insertion (Sup Fig 3B).

All-by-all genome alignment across the 26 lines revealed a total of ∼19 million pairwise structural variants (Sup Table 1 and 2), among which 22.5% were simple deletions or insertions and 77.5% showed a reciprocal unalignment between reference and query (Sup Fig 2). We also identified 5314 large tandem duplications (>10 kb), including segmental duplications (Sup Fig 4A) and nested duplications (Sup Fig 4B). Many large inversions and duplications have sustained additional insertions and rearrangements, suggesting ancient origins.

### Pan-genome analysis

#### Pangenome space

We employed the all-by-all syntenic alignments to calculate the pan-genome space (Sup Fig 5A). The added non-redundant genome size was calculated upon the addition of each genome, which was subsequently used as the reference to investigate the expanded genomic space. Aligned segments between the *n*th genome and all its predecessor genomes (*n-1*) were merged, and unaligned segments of the *n*th genome were retained. The unaligned parts of each additional line are the novel regions added to the pan-genome space. The order of NAM lines was shuffled 1000 times, and pan-genome was calculated for every case. The pipeline for pangenome computation and permutation was implemented in script pangenome_cal.py.

#### Frequency of B73-like genome space in NAM population

The frequency distribution of B73-like genomic sequences (Sup Fig 5B) was calculated by quantifying the presence/absence of every locus among 25 NAM lines. The start and end coordinates of each unit are intervals between adjacent alignment breakpoints of B73. For each chromosome, the SV breakpoints were extracted and sorted by position, and adjacent breakpoints smaller than 20bp were merged as their midpoint. Intervals between breakpoints were derived, and the occurrences of each interval were counted across NAM based on genome alignment. B73 segments present in 25 lines were represented by an allele frequency of 26, and an allele frequency of 1 depicts B73-specific regions. The above steps were conducted with script allele_frequency_cal.py. Genes and UMRs that overlap with each interval were identified with bedtools intersect, and subsequently quantified for every allele frequency.

### Structural comparison among repeat arrays

To measure the genetic distance of syntenic repeat arrays, we employed the dot-matrix method to perform pairwise sequence alignment for repeat arrays, and calculated pairwise distance based on the number of monomer matches between reference and query. This pipeline was carried out for the structural comparison of ten CentC arrays, ten classical knobs and the nucleolus organizer region (NOR) across 26 NAM lines. As large knobs are intermingled with knob180 and TR-1 monomers (Liu et al. 2020; Hufford et al. 2021), we performed alignment with the dominant monomer type in the array.

#### Syntenic repeat arrays identification

The coordinates of syntenic knobs, CentC, and NOR arrays were obtained from a prior study (Hufford et al. 2021). We identified CentC arrays located within 5 Mb upstream and downstream of the active centromeres (defined by CENH3 ChIP-seq; (Wang et al. 2021)) for each chromosome as true centromeric arrays. Classical knobs located on 2L, 3L, 4L, 5L, 6S1, 6S2, 6L, 7L, 8L, 9S were selected for analysis. The NOR regions are present in syntenic areas on the short arm of chromosome 6 in all lines.

#### Pairwise alignment via dot-matrix

As minimap2 failed to align tandem repetitive areas, we employed the dot-matrix approach to perform pairwise alignments between repeat arrays (Gibbs and McIntyre 1970). The traditional dot-matrix method compares two sequences through identifying nucleotide or amino acid matches on the main diagonal. In our pipeline, repeat arrays from reference and query were regarded as two sequences, where each repeat monomer was analyzed as a single residue. To identify the monomer pairs that share a common ancestor, we aligned all monomers from the reference array to those from the query array and measured their genetic distance. A match was assigned to a monomer pair when their similarity exceeded a certain threshold, and a dot was placed in the matrix. Structural similarity between the two repeat arrays was evaluated through manually inspecting the main diagonal in the dot-matrix.

To construct the dot matrix, monomer indexes from reference and query arrays were written along the two axes, where *n* represents the *n*th monomer from each array. Sequences of indexed monomers were extracted with bedtools getfasta (v2.29.2; -nameOnly -s). Genetic distance between any monomer pairs was measured through all-by-all alignments (*i* x *j*) with BLAT (v3.5;-minIdentity=70 -maxGap=10 -minScore=0 -repMatch=2147483647). The similarity score for each monomer alignment was calculated with Jaccard Index: *Len*(*A, B*) / {*Len*(*A*) + *Len*(*B*) − *Len*(*A, B*)}, where *Len*(*A*) and *Len*(*B*) represent the lengths of monomers A and B, and *Len*(*A, B*) is the number of matched nucleotides between them. Monomer pairs with a jaccard index above 0.98 were classified as matches and marked in the matrix. Dot matrices were plotted with R and structural similarity evaluated manually.

#### Similarity calculation and clustering

To assess the overall similarity between two repeat arrays in a quantitative manner, we measured the total number of monomer matches for each alignment, and normalized it against the length of the smaller array to account for the difference in array length. The similarity of each pair of repeat arrays among NAM was calculated and used as input to construct a correlation matrix. This correlation matrix was visualized as a network through qgraph in R.

### Divergence-time estimation with the whole-genome alignment method

#### Whole chromosome intergenic spaces

As the syntenic anchors represent the true common ancestry between reference and query genomes, SNPs located in these syntenic regions could be used to accurately infer the divergence time for individual aligned segments. The syntenic aligned segments in a minimap2 paf format were identified based on coordinates derived from the synteny identification step and subjected to variant calling with paftools.js call (v2.17) using parameters: -L50 -q0 -l50.

To obtain a more accurate estimation of divergence time, we excluded genes and unmethylated regions in each syntenic alignment (annotated in (Hufford et al. 2021)) and counted SNPs in remaining areas, which are true intergenic spaces. Accordingly, the effective alignment length for each region was calculated as follows: *len*(*syntenic aligned region*) − *len*(*union*(*genes* + *UMRs*)). Divergence time for each aligned segment was further estimated with equation *d*/*u*/2, where *d* is the total number of SNPs over individual effective alignment length, and *u* is the molecular clock of 3.3 × 10^−8 (Clark et al. 2005)^.

The reference genome was divided into 20 kb non-overlapping windows and syntenic SNPs were projected to each window with bedtools (v2.29.2). The divergence time was calculated with a molecular clock of 3.3 × 10^−8^ by summarizing the total SNPs against an effective alignment length in window: 20,000 − *sum*(*len*(*syntenic unaligned regions*)) − *sum*(*len*(*union*(*genes* + *UMRs*))). We also tested a broader range of window sizes, including 10 kb, 30 kb, 40 kb, 50 kb and 100 kb to estimate the effect of window size on divergence time profiling (Sup Fig 15).

We also profiled divergence time distribution over cenhaps only with a window size of 20 kb in density plots. The B73 coordinates of cenhap regions are as follows: chr1 (134 to 140 Mb), chr2 (96 to 101 Mb), chr3 (80 to 90 Mb), chr4 (102 to 112 Mb), chr5 (103 to 110 Mb), chr6 (57 to 63 Mb), chr7 (60 to 70 Mb), chr8 (46 to 51 Mb), chr9 (56 to 63 Mb), chr10 (42 to 50 Mb).

#### Knobs

As knobs are interspersed with a variety of transposable elements (Liu et al. 2020; Hufford et al. 2021), large portions of syntenic knobs could be uniquely aligned -- mostly over syntenic TEs but also including many repeat monomers. To eliminate the effect of incorrect chaining of repeat sequences, we removed SNPs in TR-1 and knob180 repeat sequences with bedtools intersect (v2.29.2) and computed the divergence time of each aligned segment with SNPs located in non-tandem repetitive areas.

We were not able to estimate the times of divergence for CentC arrays by this method because most embedded TEs are of the CRM class and show high homology to each other, creating erroneous alignments. Divergence times for cenhaps were estimated with the selected pericentromeric regions as described above.

### Hierarchical clustering over pericentromeric regions

To investigate the relative evolutionary distance among cenhaps, we used syntenic SNPs for clustering analysis. These SNPs were obtained through the divergence time estimation step described above. The B73 coordinates of cenhap regions are as follows: chr1 (134 to 140 Mb), chr2 (96 to 101 Mb), chr3 (80 to 90 Mb), chr4 (102 to 112 Mb), chr5 (103 to 110 Mb), chr6 (57 to 63 Mb), chr7 (60 to 70 Mb), chr8 (46 to 51 Mb), chr9 (56 to 63 Mb), chr10 (42 to 50 Mb). All SNPs projected to B73 in each cenhap interval were cataloged and used as input for distance matrix construction. This matrix has a dimension of *NXM*, where *N* represents 26 NAM lines, and *M* is the total number of positions in B73 where SNPs were identified between any line and the reference genome. The absence of SNPs in the matrix could be accounted for by 1) high sequence divergence, where regions can not be aligned to call SNPs, or 2) high sequence similarity, in which case no SNPs are found over aligned areas. To differentiate the two causes, we marked unaligned segments in the matrix as NA. Pericentromeric SNPs that are shared across all lines were subjected to hierarchical clustering with scikit-learn.

### Divergence-time estimation with short-read mapping

#### Source of data

Paired-end Illumina data of 49 *parviglumis* lines from Palmar Chico in Balsas river drainage of Mexico were obtained from Bioproject PRJNA616247 (SRR11448786-SRR11448838). Illumina reads of 14 *parviglumis* lines TIL01 (SRR447882), TIL02 (SRR447886), TIL03 (SRR447894-SRR447895), TIL04 (SRR447962-SRR447964), TIL05 (SRR447755-SRR447757),TIL06 (SRR447827-SRR447829), TIL07(SRR447960-SRR447961), TIL09 (SRR447954-SRR447955), TIL10 (SRR447825-SRR447826), TIL11 (SRR5976511), TIL12 (SRR447997), TIL14 (SRR447780-SRR447782), TIL15 (SRR447859-SRR447860) and TIL17 (SRR447896-SRR447898) were from HapMap II project SRP011907. Paired-end reads of two *mexicana* lines, TIL08 (SRR447933-SRR447934) and TIL25 (SRR447936-SRR5976310), were obtained from study SRP011907, and data from two other *mexicana* lines (SRR7758236 and SRR7758237) were downloaded from project PRJNA487810. Reads for *Zea mays huehuetenangensis* samples Hue2 and Hue4 were downloaded from PRJNA384363, and data from *Zea diploperennis* (SRR13687522) were from project PRJNA700589. Paired-end data for two *Tripsacum dactyloides* (sister genus of *Zea*) lines TDD39103 (SRR447804-SRR447807) and Trip_ISU_1 (SRR7758238) were from SRP011907 and PRJNA487810. Short-read data from the NAM lines were obtained from PRJEB31061 and PRJEB32225.

#### SNP calling

Illumina reads were trimmed with trimgalore (v0.6.5) and aligned to B73 RefGen_v5 with bwa-mem (v0.7.17). Variant calling was conducted on bam files with a mapping quality above 20 using bcftools mpileup (v1.6; -Ou -f -C50). To reduce the effect of read mapping errors on SNP calling, sites with too low or too high read depth were removed using bcftools filter, where the lower and upper bounds were respectively defined as ¼ and 4 times of the mean read depth of input bam files. High-confidence calls were obtained by further applying a quality cutoff of 20, and homozygous SNPs were extracted with bcftools view.

#### Divergence calculation and normalization

To estimate the divergence of each line from the reference with short reads, we calculated the genetic distance between each sample and reference in a fixed window. The reference genomes were divided into 20 kb non-overlapping windows with bedtools (v2.29.2). Genetic distance was measured as the proportion of intergenic SNPs over effective SNPable lengths in each window. We applied the same coverage cutoff used for SNP calling to estimate effective length, which is between ¼ and 4 times of the mean read depth. The portion of SNPable segments that overlap with genes and UMRs were removed with bedtools (v2.29.2) to account for intergenic regions. Divergence time over each window was estimated with *d/2/u*, where u = 3.3×10^−8^.

### MSMC (Multiple Sequentially Markovian Coalescent) analysis

To infer the dynamics of maize effective population size over the past million years, we employed the MSMC method to analyze syntenic intergenic SNPs between the 25 NAM lines and B73. As low residual heterozygosity was identified among NAM inbreds, one haplotype was used for each line for MSMC analysis. To generate input files for msmc2 (Schiffels and Durbin 2014; Schiffels and Wang 2020), SNPs in VCF format across 25 lines were merged with bcftools (v1.6; http://samtools.github.io/bcftools) and phased with BEAGLE (Browning et al.). Syntenic aligned regions derived from the whole genome alignment and SV calling section between NAM and B73 were used as mappability masks to define high-confidence mapping. Phased haplotypes and mask files for 25 lines were concatenated into a single file with generate_multihetsep.py, and used as input for msmc2 (Schiffels and Wang 2020). The time segment patterning parameter was set as 5*4+25*2+5*4 for msmc2 analysis and a molecular clock of 3.3×10^−8^ was used for time estimation.

## Data Availability

The code used in this study is available at Github repository: https://github.com/jl03308/Archaic-Introgression. SV metadata (left and right breakpoint coordinates of each pairwise comparison in text file format) are in the github SV-calling section.

## ACKNOWLEDGEMENTS

We thank James Leebens-Mack who provided encouragement, comments, and edits to the final manuscript. We also thank Matthew Hufford, Jonathan Gent, Michelle Stitzer, David Wills, Meghan Brady and Rebecca Piri for comments to earlier versions of the manuscript. This work was supported by grants from the National Science Foundation (IOS-174400 and IOS-2040218).

## Notes

### Competing Interest Statement

The authors have declared no competing interest.

## References

Abouelhoda MI, Ohlebusch E. 2005. Chaining algorithms for multiple genome comparison. J Discrete Algorithms 3: 321–341.

Albert PS, Gao Z, Danilova TV, Birchler JA. 2010. Diversity of chromosomal karyotypes in maize and its relatives. Cytogenet Genome Res 129: 6–16.

Altemose N, Logsdon GA, Bzikadze AV, Sidhwani P, Langley SA, Caldas GV, Hoyt SJ, Uralsky L, Ryabov FD, Shew CJ, et al. 2021. Complete genomic and epigenetic maps of human centromeres. bioRxiv 2021.07.12.452052. https://www.biorxiv.org/content/10.1101/2021.07.12.452052v2.external-links.html (Accessed December 13, 2021).

Andorf C, Beavis WD, Hufford M, Smith S, Suza WP, Wang K, Woodhouse M, Yu J, Lübberstedt T. 2019. Technological advances in maize breeding: past, present and future. Theor Appl Genet 132: 817–849.

Ardelean CF, Becerra-Valdivia L, Pedersen MW, Schwenninger J-L, Oviatt CG, Macías-Quintero JI, Arroyo-Cabrales J, Sikora M, Ocampo-Díaz YZE, Rubio-Cisneros II, et al. 2020. Evidence of human occupation in Mexico around the Last Glacial Maximum. Nature 584: 87–92.

Becerra-Valdivia L, Higham T. 2020. The timing and effect of the earliest human arrivals in North America. Nature 584: 93–97.

Beissinger TM, Wang L, Crosby K, Durvasula A, Hufford MB, Ross-Ibarra J. 2016. Recent demography drives changes in linked selection across the maize genome. Nat Plants 2: 16084.

Browning BL, Zhou Y, Browning SR. A one penny imputed genome from next generation reference panels. http://dx.doi.org/10.1101/357806.

Brunner S, Fengler K, Morgante M, Tingey S, Rafalski A. 2005. Evolution of DNA sequence nonhomologies among maize inbreds. Plant Cell 17: 343–360.

Calfee E, Gates D, Lorant A, Taylor Perkins M, Coop G, Ross-Ibarra J. 2021. Selective sorting of ancestral introgression in maize and teosinte along an elevational cline. bioRxiv 2021.03.05.434040. https://www.biorxiv.org/content/10.1101/2021.03.05.434040v3 (Accessed October 26, 2021).

Chia J-M, Song C, Bradbury PJ, Costich D, de Leon N, Doebley J, Elshire RJ, Gaut B, Geller L, Glaubitz JC, et al. 2012. Maize HapMap2 identifies extant variation from a genome in flux. Nat Genet 44: 803–807.

Choi K, Zhao X, Tock AJ, Lambing C, Underwood CJ, Hardcastle TJ, Serra H, Kim J, Cho HS, Kim J, et al. 2018. Nucleosomes and DNA methylation shape meiotic DSB frequency in Arabidopsis thaliana transposons and gene regulatory regions. Genome Res 28: 532–546.

Clark RM, Linton E, Messing J, Doebley JF. 2004. Pattern of diversity in the genomic region near the maize domestication gene tb1. Proc Natl Acad Sci U S A 101: 700–707.

Clark RM, Tavaré S, Doebley J. 2005. Estimating a nucleotide substitution rate for maize from polymorphism at a major domestication locus. Mol Biol Evol 22: 2304–2312.

Dawe RK, Lowry EG, Gent JI, Stitzer MC, Swentowsky KW, Higgins DM, Ross-Ibarra J, Wallace JG, Kanizay LB, Alabady M, et al. 2018. A Kinesin-14 Motor Activates Neocentromeres to Promote Meiotic Drive in Maize. Cell 173: 839–850.e18.

Doebley J. 2004. The genetics of maize evolution. Annu Rev Genet 38: 37–59.

Finseth FR, Nelson TC, Fishman L. 2021. Selfish chromosomal drive shapes recent centromeric histone evolution in monkeyflowers. PLoS Genet 17: e1009418.

Fu H, Dooner HK. 2002. Intraspecific violation of genetic colinearity and its implications in maize. Proc Natl Acad Sci U S A 99: 9573–9578.

Gent JI, Wang N, Dawe RK. 2017. Stable centromere positioning in diverse sequence contexts of complex and satellite centromeres of maize and wild relatives. Genome Biol 18: 121.

Ghaffari R, Cannon EKS, Kanizay LB, Lawrence CJ, Dawe RK. 2013. Maize chromosomal knobs are located in gene-dense areas and suppress local recombination. Chromosoma 122: 67–75.

Gibbs AJ, McIntyre GA. 1970. The diagram, a method for comparing sequences. Its use with amino acid and nucleotide sequences. Eur J Biochem 16: 1–11.

Gore MA, Chia J-M, Elshire RJ, Sun Q, Ersoz ES, Hurwitz BL, Peiffer JA, McMullen MD, Grills GS, Ross-Ibarra J, et al. 2009. A first-generation haplotype map of maize. Science 326: 1115–1117.

Haberer G, Kamal N, Bauer E, Gundlach H, Fischer I, Seidel MA, Spannagl M, Marcon C, Ruban A, Urbany C, et al. 2020. European maize genomes highlight intraspecies variation in repeat and gene content. Nat Genet 52: 950–957.

Hilton H, Gaut BS. 1998. Speciation and domestication in maize and its wild relatives: evidence from the globulin-1 gene. Genetics 150: 863–872.

Huang Y, Huang W, Meng Z, Braz GT, Li Y, Wang K, Wang H, Lai J, Jiang J, Dong Z, et al. 2021. Megabase-scale presence-absence variation with Tripsacum origin was under selection during maize domestication and adaptation. Genome Biol 22: 237.

Hufford MB, Lubinksy P, Pyhäjärvi T, Devengenzo MT, Ellstrand NC, Ross-Ibarra J. 2013. The genomic signature of crop-wild introgression in maize. PLoS Genet 9: e1003477.

Hufford MB, Seetharam AS, Woodhouse MR. 2021. De novo assembly, annotation, and comparative analysis of 26 diverse maize genomes. bioRxiv. https://www.biorxiv.org/content/10.1101/2021.01.14.426684v1.abstract.

Hufford MB, Xu X, van Heerwaarden J, Pyhäjärvi T, Chia J-M, Cartwright RA, Elshire RJ, Glaubitz JC, Guill KE, Kaeppler SM, et al. 2012. Comparative population genomics of maize domestication and improvement. Nat Genet 44: 808–811.

Lahn BT, Page DC. 1999. Four evolutionary strata on the human X chromosome. Science 286: 964–967.

Lampson MA, Black BE. 2017. Cellular and Molecular Mechanisms of Centromere Drive. Cold Spring Harb Symp Quant Biol 82: 249–257.

Langley SA, Miga KH, Karpen GH, Langley CH. 2019. Haplotypes spanning centromeric regions reveal persistence of large blocks of archaic DNA. Elife 8. http://dx.doi.org/10.7554/eLife.42989.

Leyden BW, Brenner M, Hodell DA, Curtis JH. 2013. Late Pleistocene climate in the central American lowlands. In Climate Change in Continental Isotopic Records, pp. 165–178, American Geophysical Union, Washington, D. C.

Li H. 2018. Minimap2: pairwise alignment for nucleotide sequences. Bioinformatics 34: 3094– 3100.

Liu J, Seetharam AS, Chougule K, Ou S, Swentowsky KW, Gent JI, Llaca V, Woodhouse MR, Manchanda N, Presting GG, et al. 2020. Gapless assembly of maize chromosomes using long-read technologies. Genome Biol 21: 121.

Liu S, Yeh C-T, Ji T, Ying K, Wu H, Tang HM, Fu Y, Nettleton D, Schnable PS. 2009. Mu transposon insertion sites and meiotic recombination events co-localize with epigenetic marks for open chromatin across the maize genome. PLoS Genet 5: e1000733.

Lynch M, Ackerman MS, Gout J-F, Long H, Sung W, Thomas WK, Foster PL. 2016. Genetic drift, selection and the evolution of the mutation rate. Nat Rev Genet 17: 704–714.

Malik HS. 2009. The centromere-drive hypothesis: a simple basis for centromere complexity. Prog Mol Subcell Biol 48: 33–52.

Matsuoka Y, Vigouroux Y, Goodman MM, Sanchez G J, Buckler E, Doebley J. 2002. A single domestication for maize shown by multilocus microsatellite genotyping. Proc Natl Acad Sci U S A 99: 6080–6084.

McMullen MD, Kresovich S, Villeda HS, Bradbury P, Li H, Sun Q, Flint-Garcia S, Thornsberry J, Acharya C, Bottoms C, et al. 2009. Genetic properties of the maize nested association mapping population. Science 325: 737–740.

Monroe JG, Srikant T, Carbonell-Bejerano P, Becker C, Lensink M, Exposito-Alonso M, Klein M, Hildebrandt J, Neumann M, Kliebenstein D, et al. 2022. Mutation bias reflects natural selection in Arabidopsis thaliana. Nature. http://dx.doi.org/10.1038/s41586-021-04269-6.

Nambiar M, Smith GR. 2016. Repression of harmful meiotic recombination in centromeric regions. Semin Cell Dev Biol 54: 188–197.

Nordborg M. 2004. Coalescent Theory. In Handbook of Statistical Genetics, John Wiley & Sons, Ltd, Chichester.

Patton AH, Margres MJ, Stahlke AR, Hendricks S, Lewallen K, Hamede RK, Ruiz-Aravena M, Ryder O, McCallum HI, Jones ME, et al. 2019. Contemporary Demographic Reconstruction Methods Are Robust to Genome Assembly Quality: A Case Study in Tasmanian Devils. Mol Biol Evol 36: 2906–2921.

Piperno DR, Ranere AJ, Holst I, Iriarte J, Dickau R. 2009. Starch grain and phytolith evidence for early ninth millennium B.P. maize from the Central Balsas River Valley, Mexico. Proc Natl Acad Sci U S A 106: 5019–5024.

Rani S, Rajpoot DS. 2016. LIS using backtracking and branch-and-bound approaches. CSI Transactions on ICT 4: 87–93.

Ross-Ibarra J, Tenaillon M, Gaut BS. 2009. Historical divergence and gene flow in the genus Zea. Genetics 181: 1399–1413.

Sanmiguel P, Bennetzen JL. 1998. Evidence that a Recent Increase in Maize Genome Size was Caused by the Massive Amplification of Intergene Retrotransposons. Ann Bot 82: 37–44.

Schiffels S, Durbin R. 2014. Inferring human population size and separation history from multiple genome sequences. Nat Genet 46: 919–925.

Schiffels S, Wang K. 2020. MSMC and MSMC2: The Multiple Sequentially Markovian Coalescent. Methods Mol Biol 2090: 147–166.

Schneider KL, Xie Z, Wolfgruber TK, Presting GG. 2016. Inbreeding drives maize centromere evolution. Proc Natl Acad Sci U S A 113: E987–96.

Shi J, Wolf SE, Burke JM, Presting GG, Ross-Ibarra J, Dawe RK. 2010. Widespread gene conversion in centromere cores. PLoS Biol 8: e1000327.

Song B, Marco-Sola S, Moreto M, Johnson L, Buckler ES, Stitzer MC. 2022. AnchorWave: Sensitive alignment of genomes with high sequence diversity, extensive structural polymorphism, and whole-genome duplication. Proc Natl Acad Sci U S A 119. http://dx.doi.org/10.1073/pnas.2113075119.

Sun S, Zhou Y, Chen J, Shi J, Zhao H, Zhao H, Song W, Zhang M, Cui Y, Dong X, et al. 2018. Extensive intraspecific gene order and gene structural variations between Mo17 and other maize genomes. Nat Genet 50: 1289–1295.

Swentowsky KW, Gent JI, Lowry EG, Schubert V, Ran X, Tseng K-F, Harkess AE, Qiu W, Dawe RK. 2020. Distinct kinesin motors drive two types of maize neocentromeres. Genes Dev 34: 1239–1251.

Tenaillon MI, U’Ren J, Tenaillon O, Gaut BS. 2004. Selection versus demography: a multilocus investigation of the domestication process in maize. Mol Biol Evol 21: 1214–1225.

Tittes S, Lorant A, McGinty S, Doebley JF, Holland JB, de Jesus Sánchez-González J, Seetharam A, Tenaillon M, Ross-Ibarra J. 2021. Not so local: the population genetics of convergent adaptation in maize and teosinte. bioRxiv 2021.09.09.459637. https://www.biorxiv.org/content/10.1101/2021.09.09.459637v1.full.pdf+html (Accessed October 12, 2021).

van Heerwaarden J, Doebley J, Briggs WH, Glaubitz JC, Goodman MM, de Jesus Sanchez Gonzalez J, Ross-Ibarra J. 2011. Genetic signals of origin, spread, and introgression in a large sample of maize landraces. Proc Natl Acad Sci U S A 108: 1088–1092.

Wang L, Beissinger TM, Lorant A, Ross-Ibarra C, Ross-Ibarra J, Hufford MB. 2017. The interplay of demography and selection during maize domestication and expansion. Genome Biol 18: 215.

Wang N, Liu J, Ricci WA, Gent JI, Dawe RK. 2021. Maize centromeric chromatin scales with changes in genome size. Genetics 217: iyab020

Williams JW. 2003. Variations in tree cover in North America since the last glacial maximum. Glob Planet Change 35: 1–23.

Wright S. 1949. The genetical structure of populations. Ann Eugen 15: 323–354.

